# Detection of diadromous fish using environmental DNA: prospects for its use in conservation of endangered species occurring in Portugal

**DOI:** 10.1101/2025.03.27.645525

**Authors:** Sofia Duarte, Ana Matilde Santos, Carlos Antunes, Ronaldo Sousa, Filipe O. Costa

## Abstract

The high economic value and cultural relevance of diadromous fish make them primary targets for traditional fisheries, that need effective management to ensure the long-term survival of their populations. In Portugal, these species are experiencing a marked decline, primarily due to habitat loss and fragmentation, alongside pollution and overfishing. Accurate monitoring is therefore essential to strengthen the management of diadromous fish within Portuguese ecosystems. Advancements in monitoring methodologies could benefit from more sensitive approaches, such as environmental DNA (eDNA) analysis, which is increasingly recognized as a valuable complement to conventional fish monitoring. However, eDNA-based tools remain largely absent and untested in Portugal’s monitoring and management of diadromous fish. This study reviews literature on eDNA-based detection of diadromous fish species and discusses key methodological aspects influencing the detection efficiency, including sample processing (e.g., water filtration), DNA extraction methods, marker regions and primers, approach and platforms. A particular focus is placed on diadromous fishes occurring in Portugal, which includes several endangered species, and the prospects of using eDNA to monitor them. We also report a pilot study comparing specific assays and DNA metabarcoding in eDNA-based detection of sea lamprey in the Minho River watershed. By consolidating current literature, this work underscores the potential of eDNA to strengthen diadromous fish conservation in Portugal and offers insights to support the integration of eDNA-based tools into national monitoring frameworks. Although our study focuses on Portuguese species, the approaches and insights discussed are broadly applicable and can inform conservation efforts in other regions facing similar challenges.

**Graphical Abstract:** 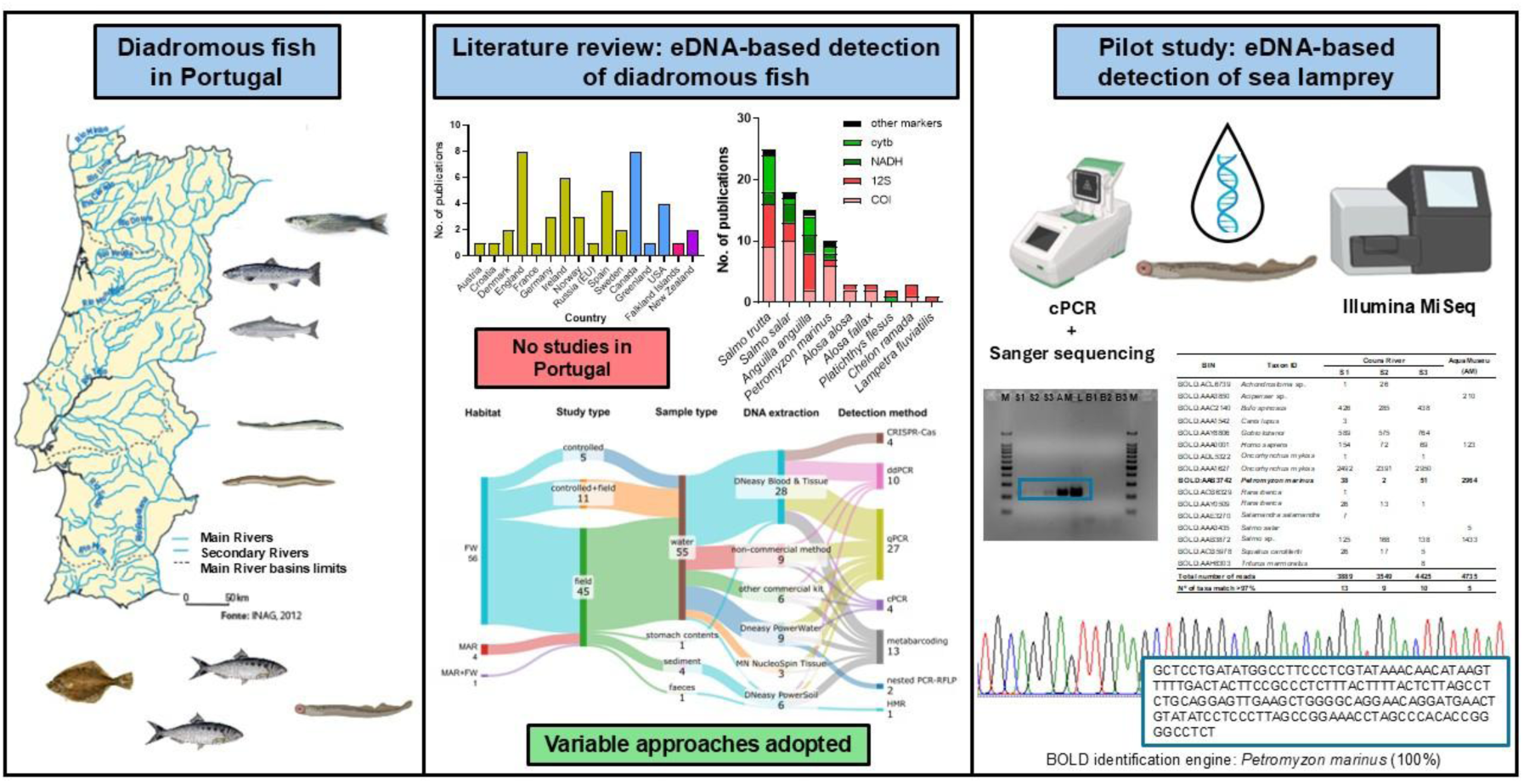

## Introduction

Diadromous fish are species characterized by regular migration between freshwater and marine environments throughout their life cycle (Jones 2006; Almeida et al. 2018). Based on their reproductive habitat, diadromous species can be further classified as catadromous, wherein reproduction occurs in the marine environment while most of their lives could be spent in freshwaters (e.g., eel), or anadromous, where spawning occur in freshwater habitats, but a significant portion of their lives is spent in the sea (e.g., shads, sea lamprey). The high commercial value attributed to diadromous fish renders them prime targets for traditional fisheries mainly in rivers, underscoring the need for meticulous management to prevent overfishing and ensure the enduring survival of their populations. Moreover, the fragmentation of river connectivity, precipitated by the construction of dams and other physical infrastructures, further exacerbates the decline in diadromous fish species populations (Mota, Micaela et al. 2016).

In Portugal, the study of diadromous fish has been ongoing for over two decades (Almeida et al. 2018). During this time, populations have steadily declined, primarily due to human activities such as habitat destruction from dam construction, which obstructs upstream migration to spawning habitats, and river flow regulation. This situation is often combined by factors such as pollution, overfishing, water scarcity, and the escalating impacts of global climate change, which are particularly pronounced in regions like the Iberian Peninsula (Mota, Micaela et al. 2016). Other probably less relevant threat may include the introduction of non-native species (e.g. European catfish predating sea lamprey, Atlantic salmon and allis shad) (Boulêtreau et al. 2020; Vejřík et al. 2024). Researchers have been primarily focused on improving population status while also striving to preserve traditional fishing practices, as many of these species hold significant commercial, historical, and cultural value (e.g., the sea lamprey in the Minho region) (Almeida et al. 2018; Braga et al. 2020). In addition, these species may also had relevant ecological effects, mainly in the past when populations were much larger. In fact, diadromous fish, migrating between freshwater and marine ecosystems, act as crucial spatial subsidies by transporting nutrients across habitats. Their movements enrich ecosystems, fueling productivity and supporting many species, including in the terrestrial realm (Schindler et al. 2003). Spawning migrations deposit marine-derived nutrients inland, benefiting freshwater food webs, while juvenile migrations enhance marine ecosystems, underscoring their ecological role as nutrient conduits.

Ten species of diadromous fish have been documented in Portugal: sea lamprey (*Petromyzon marinus* Linnaeus, 1758), European river lamprey (*Lampetra fluviatilis* Linnaeus, 1758), allis shad (*Alosa alosa* (Linnaeus, 1758)), twaite shad (*A. fallax* Lacépède, 1800), Atlantic salmon (*Salmo salar* Linnaeus, 1758), and sea trout (*Salmo trutta* Linnaeus, 1758), European eel (*Anguilla anguilla* (Linnaeus, 1758)), thin-lipped grey mullet (*Liza ramada/Chelon ramada* (Risso, 1827)), European flounder (*Platichthys flesus* (Linnaeus, 1758)) and the already regionally extinct sturgeon (*Acipenser sturio* Linnaeus, 1758) (Almeida et al. 2018).

The declining populations of diadromous species pose a persistent challenge in Conservation Biology, particularly in the detection of increasingly rare species. Addressing this issue may require intensifying sampling efforts or adopting more sensitive methods (Janosik and Johnston 2015). However, conventional approaches such as costly fisheries trawl surveys present significant drawbacks. Not only are these surveys expensive and challenging to replicate, but they also often entail traditional benthic trawling methods that can be harmful to habitats and the diverse organisms inhabiting sea or river bottoms. Conversely, the adoption of more sensitive methods, particularly those leveraging environmental DNA (eDNA), are becoming increasingly popular within aquatic ecosystems and as complementary methods to conventional fish monitoring (Janosik and Johnston 2015; Fernandez et al. 2018a; Tsuji et al. 2022). The two predominant approaches for detecting eDNA in environmental samples typically involve either single-species detection or multispecies detection (Harper et al. 2018; Bylemans et al. 2019). Single-species detection relies on employing a PCR-based technique with species-specific primers, such as conventional PCR, quantitative PCR (qPCR) or droplet digital PCR (ddPCR) (Gustavson et al. 2015; Capo et al. 2019a). This method enables the specific identification and quantification (in case of qPCR or ddPCR) of targeted species within a sample. On the other hand, multispecies detection utilizes high-throughput sequencing (HTS) following PCR amplification of eDNA using universal primers targeting short DNA barcoding regions. This approach, known as eDNA metabarcoding, allows for the simultaneous identification of multiple species present in a sample, providing a comprehensive assessment of biodiversity within an environment (Thomsen and Willerslev 2015).

The initial application of environmental DNA (eDNA) was to survey the invasive American bullfrog (*Lithobates catesbeianus* Shaw, 1802) across several ponds in France (Ficetola et al. 2008). Since then, detection assays using eDNA have been developed for numerous terrestrial and aquatic species, including native, rare, endangered, or non-native species (for a comprehensive review, see (Duarte et al. 2023)). However, to our knowledge, the utilization of eDNA-based tools for monitoring diadromous fish has been limited, and in Portugal, it is virtually non-existent. This review aims to assess the status of eDNA-based tools for monitoring diadromous fish, with reported occurrence in Portugal, and explore the potential for their implementation to enhance monitoring and conservation efforts in Portugal. For that, in a first instance, a synthesis of available data on key methodological aspects influencing eDNA detection - sample pre- processing techniques (e.g., water filtration), DNA extraction protocols, marker regions and primers, and detection platforms - was performed followed by an exploration of future perspectives for implementing eDNA-based methods to enhance knowledge on diadromous fish species in Portuguese aquatic ecosystems. By consolidating current literature, this work underscores the potential of eDNA to strengthen diadromous fish conservation efforts in Portugal and offers insights to support the integration of eDNA-based tools into national monitoring frameworks. In addition, we conducted a small pilot study to evaluate the efficiency of eDNA-based methods for detecting sea lamprey in Portuguese rivers using a conventional PCR specific assay and DNA metabarcoding (Box 1).

### BOX 1

In Portugal, the sea lamprey holds ecological, economic, and cultural importance, particularly through its role in fisheries and gastronomy (Almeida et al. 2018). However, migration barriers (e.g., dams), habitat loss, and pollution pose significant threats to its breeding populations (Almeida et al. 2018). Consequently, the species is listed as “Threatened” or “Near Threatened” on national Red Lists in most countries within OSPAR Regions I–IV (Arctic Waters; Greater Celtic Seas; Bay of Biscay and Iberian Coast) (Gustavson et al. 2015; Baltazar-Soares et al. 2022a). Conservation efforts are critical to safeguarding its survival and the associated benefits. Environmental DNA (eDNA)-based detection offers a promising method for improving monitoring and conservation of sea lamprey in Portugal. A literature review identified five specific assays developed for its detection, targeting mitochondrial genes: COI (Gustavson et al. 2015; Gingera et al. 2016), Cytb (Schloesser 2018), NADH1 (Schloesser 2018) and ATP6-ATP8 (Baltazar-Soares et al. 2022a) (Table 1). These assays have been validated in regions such as North America, Ireland, and the UK, but not in Portugal. Given the genetic uniformity of COI sequences across populations in Europe and North America (Supplementary Material 1, Table S3), a COI-based assay (Gingera et al. 2016), was tested in Portugal using three experiments (Supplementary Material 2): (1) *in vitro* testing with tissue samples; (2) *in situ* testing in aquarium water from the Aquamuseum of the Minho River; and (3) field testing in the Coura River (Minho River basin). Assay validation through conventional PCR and Sanger sequencing confirmed matches with *Petromyzon marinus* (Fig. 6, Supplementary Material 1, Table S5).

## Methods

A comprehensive literature search was conducted using the Web of Science database on January 2nd, 2024, to compile a state-of-the-art overview of eDNA-based tools for detecting and monitoring diadromous fish species with reported occurrence in Portugal. The search was limited to titles, abstracts, and keywords, encompassing the terms “eDNA” OR “environmental DNA” along with the scientific and common names of all diadromous fish species occurring in Portugal nowadays as listed in (Almeida et al. 2018) and (Mota, Micaela et al. 2016) - “Petromyzon marinus” OR “sea lamprey” OR “Lampetra fluviatilis” OR “european river lamprey” OR “Alosa alosa” OR “allis shad” OR “Alosa fallax” OR “twaite shad” OR “twait shad” OR “Salmo trutta” OR “sea trout” OR “Salmo salar” OR “atlantic salmon” OR “Anguilla anguilla” OR “european eel” OR “Liza ramada” OR “Chelon ramada” OR “thin-lipped grey mullet” OR “thin lipped grey mullet” OR “thinlip grey mullet” OR “Platichthys flesus” OR “european flounder”. This search yielded a total of 77 publications spanning from 2003 to 2023 (Supplementary Material 1, Table S1). Additionally, recognizing the significance of the journals “Environmental DNA” and “Metabarcoding and Metagenomics” within the field, supplementary searches using the same keywords were conducted, resulting in the retrieval of 70 publications published between 2019 and 2023, for the journal Environmental DNA and 18 publications published between 2017 and 2023, for the journal Metabarcoding and Metagenomics (Fig. 1, Supplementary Material 1, Table S1). This extensive search allowed to gather a comprehensive collection of literature concerning eDNA-based tools developed for targeting diadromous fish species with documented occurrences in Portugal, providing valuable insights into the current state of research in this area.

**Fig. 1.**
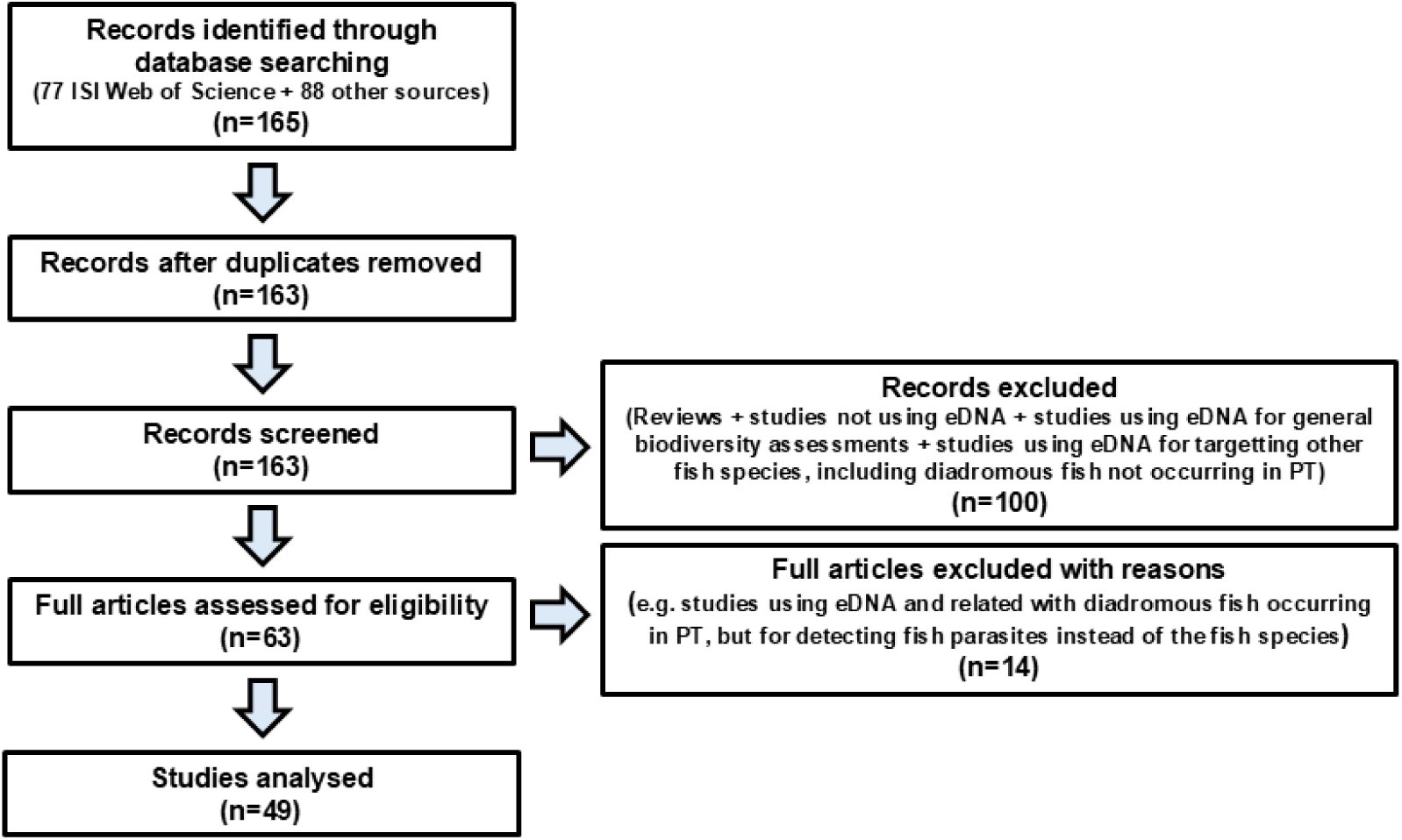
Diagram illustrating the selection process of publications included in the literature review on the use of eDNA-based methods for the detection and monitoring of diadromous fish.

Following individual inspection, 49 articles were selected for the analysis, which were published between 2015 and 2023 (Fig. 1, Supplementary Material 1, Table S2). These articles specifically addressed the utilization of eDNA-based tools for monitoring diadromous fish species with documented occurrences in Portugal. Publications that did not constitute primary research articles (e.g., reviews) or focused on species other than diadromous fish or general biodiversity assessments using DNA metabarcoding, were excluded from the analysis (Fig. 1). Although the ecotype resident brown trout live their entire lives in freshwater, unlike the anadromous sea trout, we opted to include publications targeting these populations. This decision is supported by the fact that brown trout and sea trout are ecotypes of the same species - *Salmo trutta*, and their genetic similarity poses challenges for distinguishing between the two forms using eDNA. Furthermore, both forms often coexist in the same ecosystem. For example, an analysis of 201 COI sequences downloaded from BOLD (Supplementary Material 1, Table S3) revealed no genetic differences among sequences assigned to *S. trutta*. Of these, 174 were labelled as *S. trutta*, 23 as *S. trutta fario*, 1 as *S. trutta lacustris*, and 3 as *S. trutta trutta* (sub-species not formally recognized). All sequences, and that were generated from specimens collected in 19 different countries were grouped under the same Barcode Index Number (BIN), further underscoring their genetic similarity (Supplementary Material 1, Table S3). From each selected publication, the following information was extracted: i) Geographic area/country, ii) Environment (e.g., freshwater or marine), iii) Type of environmental sample (e.g., water, sediment), iv) Whether the study was conducted in the field or in a controlled environment (e.g., aquarium, mesocosms, lab), v) Targeted species, including taxonomic classification and species category (e.g., endangered, non-native), vi) Targeted molecular markers, primers, probes (in the case of qPCR or ddPCR), and segment length (bp), vii) Platforms employed for eDNA analysis (e.g., cPCR, qPCR, DNA metabarcoding) and viii) Other relevant methodological details such as sample volume, sample pre-processing, DNA extraction methods, among others (Supplementary Material 1, Table S2). A detailed description of the methods used in the pilot study to evaluate the efficiency of eDNA-based methods for detecting sea lamprey in Portuguese rivers using specific detection and DNA metabarcoding can be found on Supplementary Material 2.

## Results and Discussion

### Status, geographic regions and habitats surveyed

An in-depth analysis of the 49 retained publications reveals a notable trend in the use of eDNA-based methodologies for detecting diadromous fish with documented occurrences in Portugal. Between 2018 and 2020, there was a significant rise in publications employing these tools (Fig. 2a). This observation aligns with the broader pattern of increasing adoption of eDNA-based techniques, particularly within aquatic ecosystems, as evidenced by recent studies (Minamoto 2022; Schenekar 2023; Duarte et al. 2023). However, this trend experienced a slight decline between 2020 and 2023, possibly due to the constraints imposed by the COVID-19 pandemic. Most of these studies were conducted in Europe (67%) (Fig. 2b), which is unsurprising given that our review focused on diadromous fish with documented occurrences in Portugal. Interestingly, no studies were conducted within Portugal itself. However, five studies were carried out in Spain, targeting three specific species (Supplementary Material 1, Table S2): European eel *Anguilla anguilla* (Burgoa Cardás et al. 2020a; Fernandez et al. 2023), Atlantic salmon *Salmo salar* (Clusa et al. 2017) and Sea/Brown trout *S. trutta* (Fernandez et al. 2018a; Fernández et al. 2019a). A smaller percentage of studies (27%) were conducted in North America, where some of these diadromous fish species are considered non-native, such as sea lamprey (Gingera et al. 2016; Schloesser 2018; Tkachuk and Dunn 2020) and brown trout (Carim et al. 2016).

**Fig. 2.**
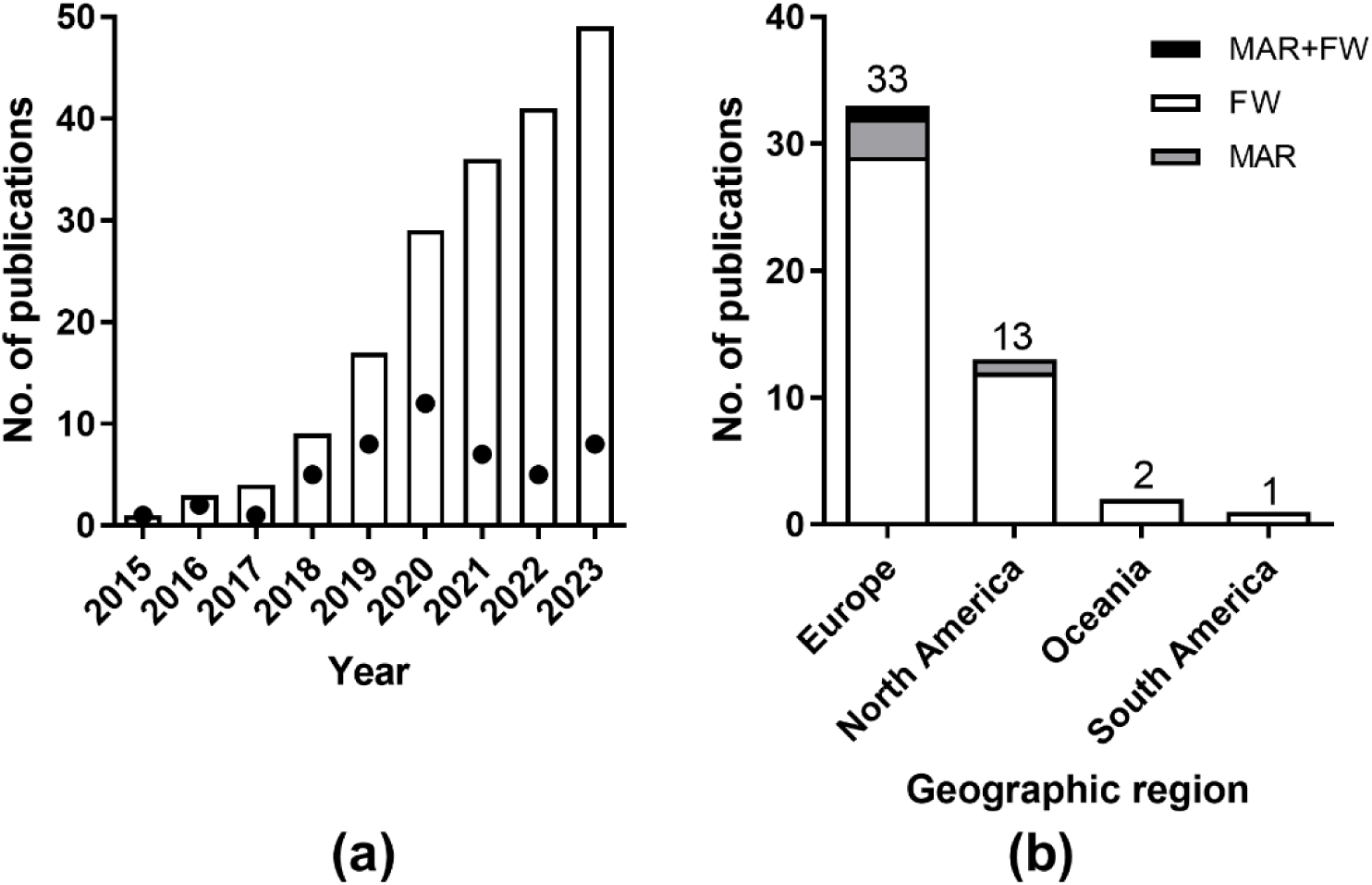
(a) Cumulative number of publications using eDNA-based tools to detect diadromous fish species with reported occurrences in Portugal. Each dot represents the number of publications identified for each year. (b) Continents and type of habitats (MAR- marine; FW – freshwater; MAR+FW – marine and freshwater) where eDNA-based tools have been employed for detecting diadromous fish species reported in Portugal.

Most studies targeted diadromous species within freshwater environments (Fig. 2, Supplementary Material 1, Table S2), although some also focused on marine environments, particularly the European eel (Jensen et al. 2018a; Barco et al. 2022; Vucić et al. 2023a) or the European flounder (Knudsen et al. 2019a) (Supplementary Material 1, Table S2), whose reproduction occurs in the marine realm. In addition, this imbalance can be attributed, in part, to the vastness, inaccessibility, and high complexity of marine environments, which pose considerable challenges for eDNA detection of rare species (Suarez-Bregua et al. 2022).

### Targeted species and most employed genetic markers in eDNA-based studies

Among diadromous fish species investigated through eDNA-based studies, *Salmo trutta* has garnered the most attention, featuring in 24 studies with 8 specific assays being developed. This prominence is likely due to the existence of two distinct ecotypes, as already above-mentioned, as well as the widespread distribution and ecological importance of the species (Gustavson et al. 2015; Carim et al. 2016; Fernandez et al. 2018a; Hernandez et al. 2020a; Laporte et al. 2020a; Halvorsen et al. 2020; Thalinger et al. 2021). It is also probable that the resident brown trout populations have been the primary focus of these studies, given their accessibility, and easier monitoring compared to the migratory sea trout (Carim et al. 2016; Minett et al. 2021a; Hallam et al. 2021). Following closely, the Atlantic salmon has been the focus of 16 studies utilizing 7 specific assays (Clusa et al. 2017; Atkinson et al. 2018a; Wood et al. 2021; Jacobsen et al. 2023), while the European eel has been studied in 15 instances and 5 specific assays have been developed for the specific detection of this species (Jensen et al. 2018a; Burgoa Cardás et al. 2020a; Halvorsen et al. 2023; Vucić et al. 2023a) (Fig. 3, Table 1, Supplementary Material 1, Table S2). In contrast, the European flounder has been the subject of only one study in the Baltic Sea (Knudsen et al. 2019a) although it has been detected in two additional studies employing DNA metabarcoding (Harper et al. 2020; Hallam et al. 2023). The thinlip grey mullet and the river lamprey have not been specifically targeted in eDNA-based studies, but have been detected in a few studies using DNA metabarcoding (Macher et al. 2021; Hallam et al. 2021, 2023). Sea lamprey has been targeted in 7 eDNA studies, with a total of 5 specific assays developed (Gustavson et al. 2015; Gingera et al. 2016; Tkachuk and Dunn 2020; Baltazar-Soares et al. 2022a). Conversely, allis shad and twaite shad have only been targeted in three studies, with only one specific assay designed, although unable to differentiate between the two species (Antognazza et al. 2019, 2021; Consuegra et al. 2021a) (Fig. 3, Table 1, Supplementary Material 1, Table S2).

**Fig. 3.**
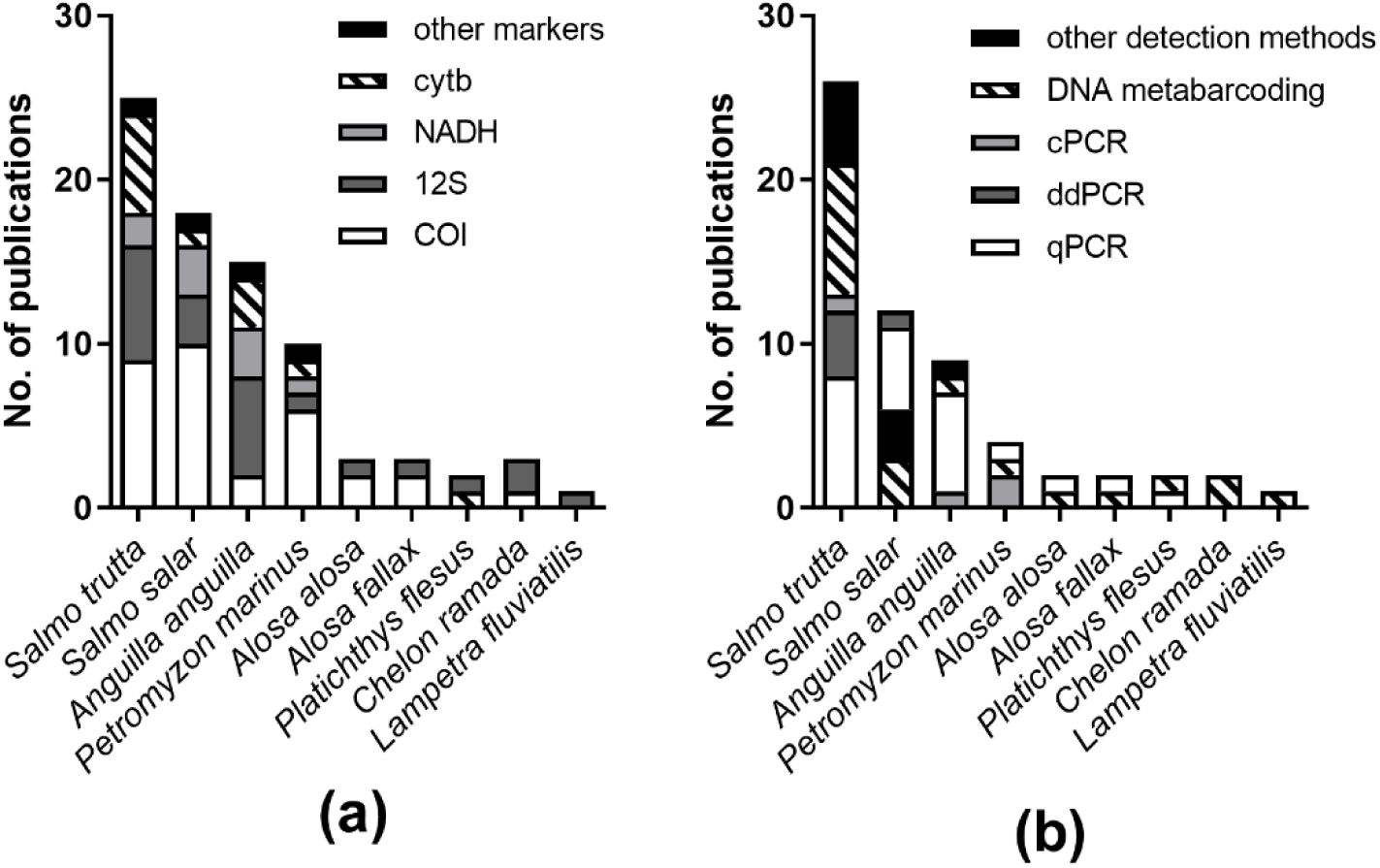
(a) Diadromous fish species with reported occurrence in Portugal and targeted in eDNA-based studies, along with the specific genetic markers used and (b) methods of detection.

**Table 1.**
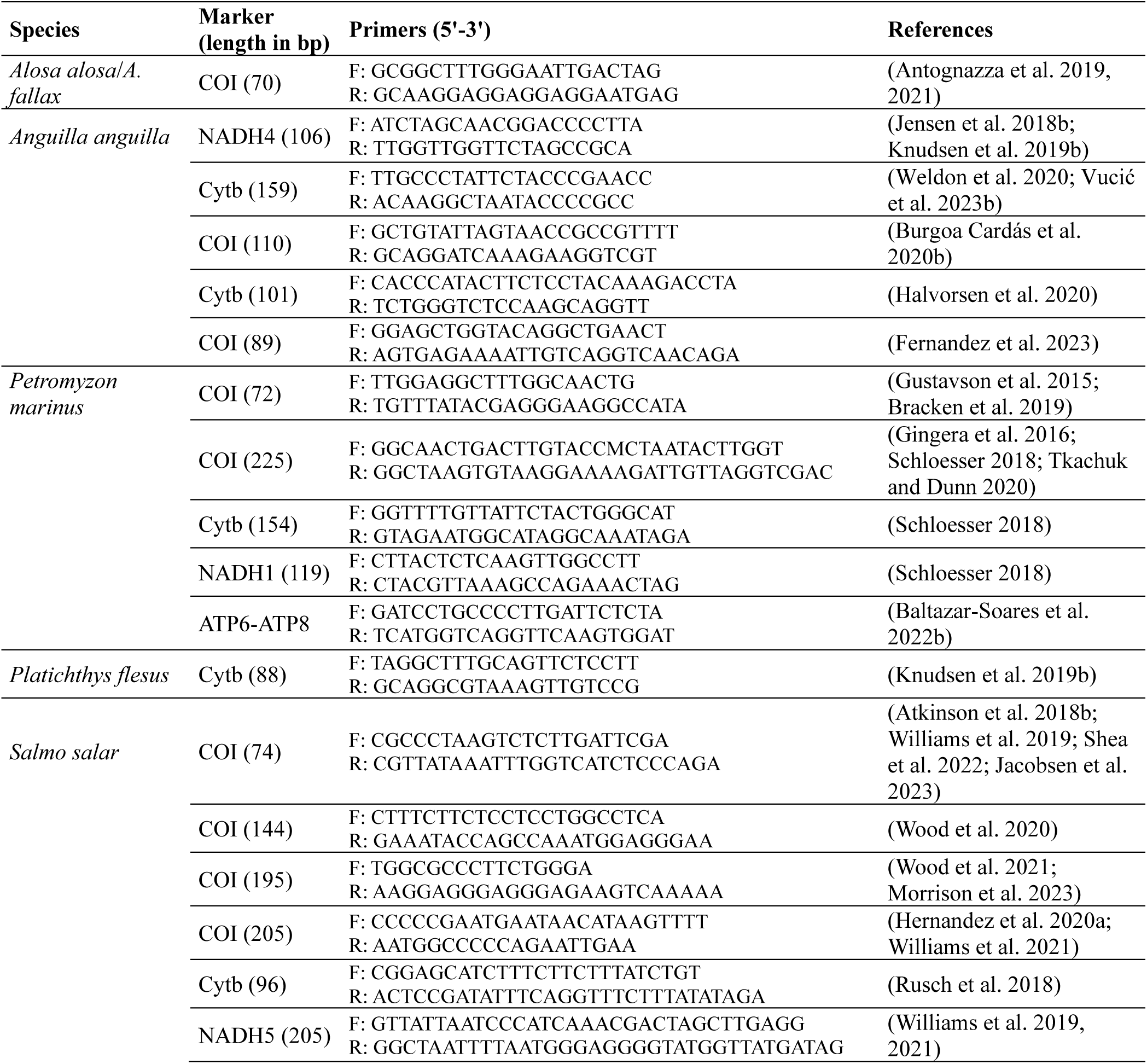
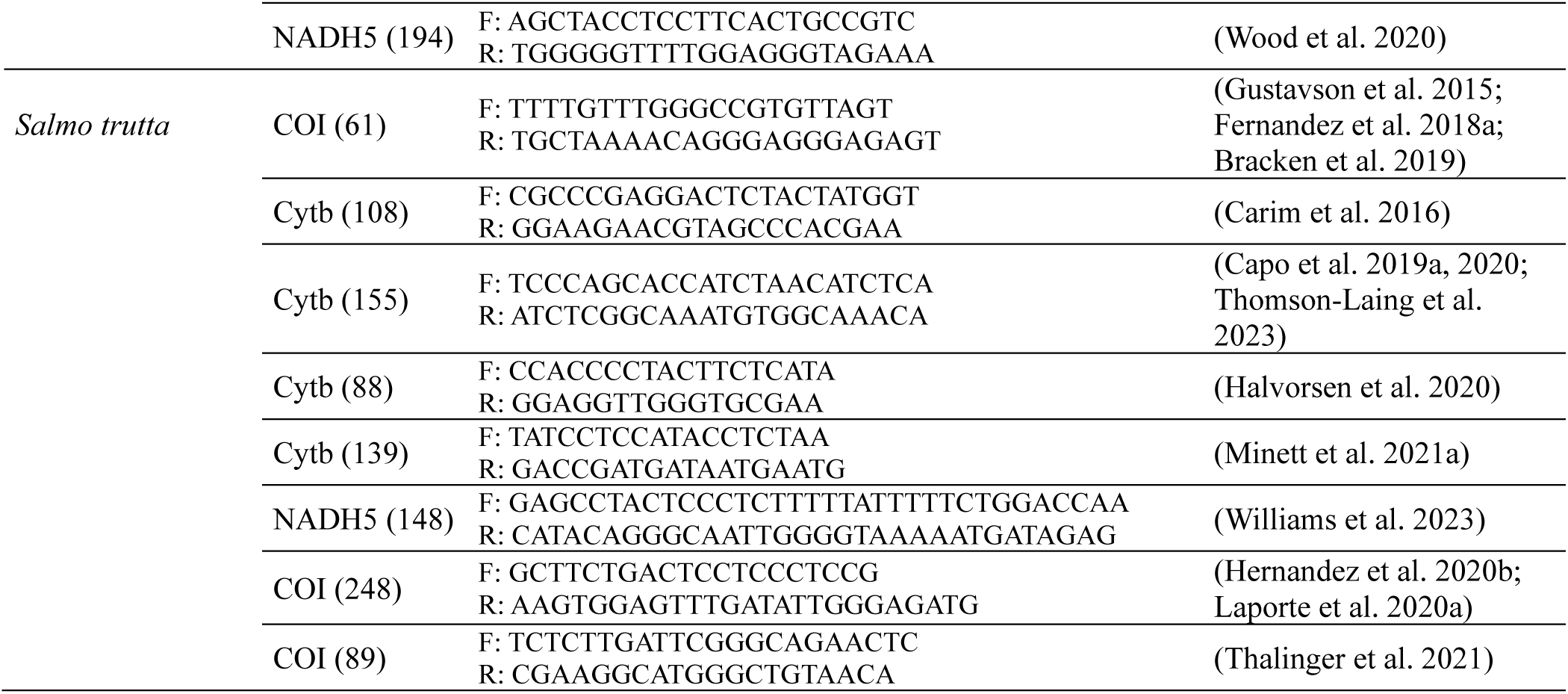
Specific assays developed for the targeted detection of diadromous fish species with reported occurrence in Portugal.

The mitochondrial cytochrome c oxidase subunit I (COI) gene has emerged as the primary target in eDNA analysis, owing to extensive characterization across species (Andújar et al. 2018). This characteristic makes it conducive to the development of specific primers and probes, facilitating precise detection (Gustavson et al. 2015; Gingera et al. 2016; Antognazza et al. 2019; Fernandez et al. 2023) (Table 1, Fig. 3a). Additionally, the cytochrome b gene (cytb) (Carim et al. 2016; Schloesser 2018; Rusch et al. 2018; Capo et al. 2019a; Halvorsen et al. 2020) and NADH genes (Jensen et al. 2018b; Halvorsen et al. 2020; Williams et al. 2021) have been frequently employed as alternative markers for diadromous fish specific detection. The high copy number of mitochondrial DNA, compared to nuclear DNA, make mitochondrial loci, such as COI, cytb and NADH genes ideal for analysis from highly degraded DNA. However, it has been argued that COI does not contain suitably conserved regions for most amplicon-based metabarcoding applications (Deagle et al. 2014). Nevertheless, the comprehensive coverage of reference sequence databases for COI (see Supplementary Material 1, Table S3), the variability inherent in this marker, and recent advancements in DNA sequencing protocols, all support the argument for standardizing the use of COI in metazoan community samples, encompassing fish as well (Andújar et al. 2018). However, the 12S rRNA gene (12S), in particular the MiFish primers (Miya et al. 2015), have been predominantly employed in DNA metabarcoding studies, offering the ability to detect multiple fish species, including the diadromous species investigated in the present study (Griffiths et al. 2020; Muha et al. 2021; Consuegra et al. 2021b; Laporte et al. 2022; Barco et al. 2022) (Supplementary Material 1, Table S2). These primers target a hypervariable region of the 12S (163-185 bp), which contains sufficient information to unambiguously identify fishes except for some closely related congeners (Miya et al. 2015).

### Methodological considerations

The sensitivity of eDNA assays is contingent upon several methodological choices, including: 1) the type of substrate sampled (sample type); 2) for water samples, factors such as the filtered volume, filter composition, pore size, and filter storage method; 3) the DNA extraction method and 4) the chosen detection methodology, which encompasses technical considerations such as the number of PCR replicates (for PCR-based methods), as well as detection thresholds (e.g., qPCR DNA metabarcoding) (Schenekar et al. 2020). The methodologies employed throughout the analytical chain of eDNA-based detection, from sampling to the detection platform, exhibited significant variability among the 49 publications targeting diadromous fish species (61 case studies in total, because some publications target more than one species, used more than one target detection method or genetic marker, among other methodological reasons) (Fig. 4, Supplementary Material 1, Table S2).

**Fig. 4.**
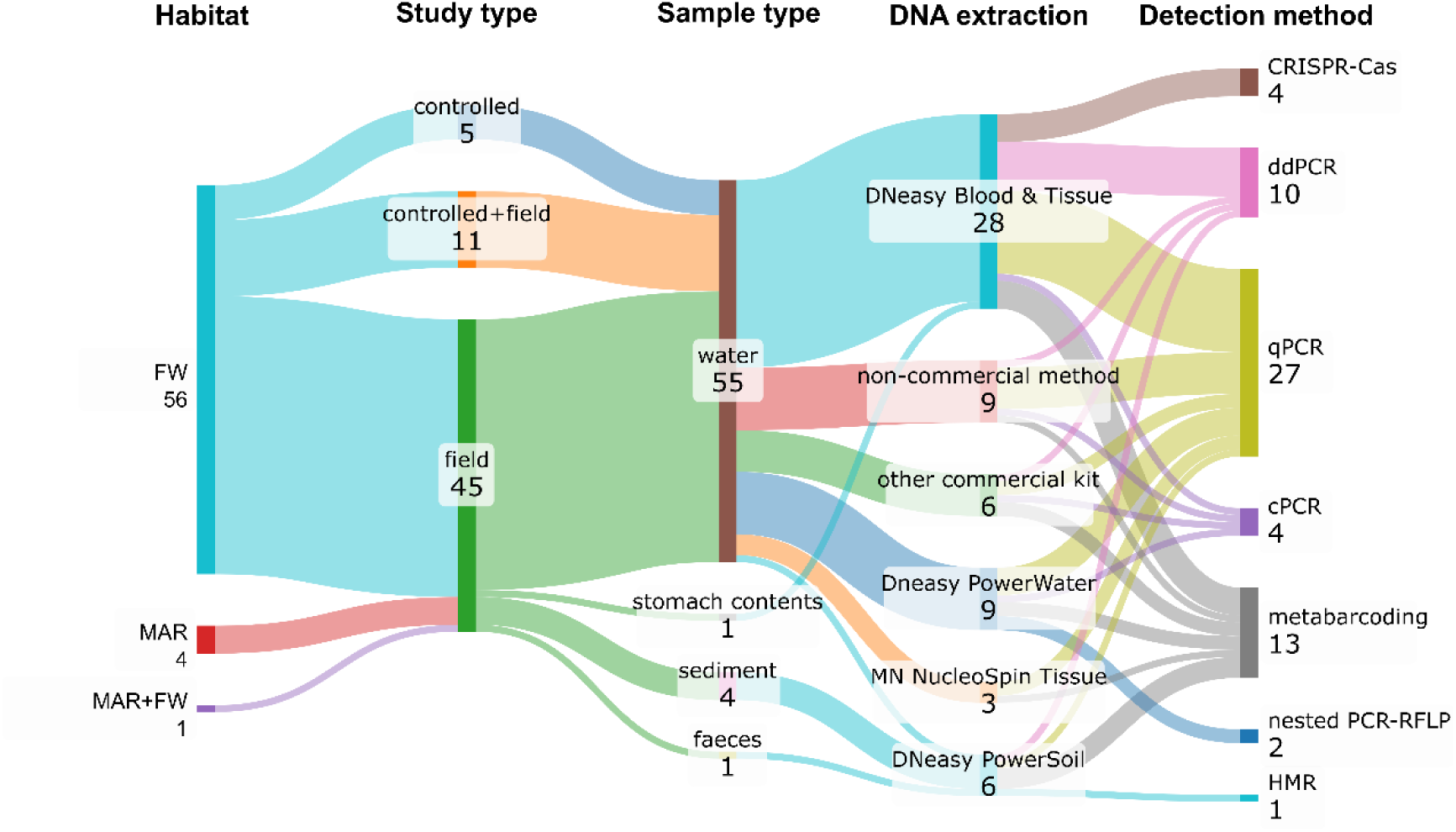
Sankey diagram resuming the most employed methodologies on each type of habitat (MAR-marine; FW – freshwater; MAR+FW – marine and freshwater) and study conducted using eDNA to detect diadromous fish species with reported occurrence in Portugal.

#### Sample type choice

Water has been the predominant sample type surveyed in eDNA-based studies targeting diadromous fish (Gustavson et al. 2015; Carim et al. 2016; Atkinson et al. 2018a; Antognazza et al. 2019; Burgoa Cardás et al. 2020a) (Supplementary Material 1, Table S2). Less commonly used substrates include sediment (Baltazar-Soares et al. 2022a; Thomson-Laing et al. 2023; Schallenberg et al. 2023), as well as stomach contents (Jensen et al. 2018a) or faeces (Harper et al. 2020) (Supplementary Material 1, Table S2), the latter two employed when investigating the presence of diadromous fish species in predator diets.

While water has undoubtedly been the most frequently sampled medium for detecting diadromous fish species through eDNA (Supplementary Material 1, Table S2, Fig. 4), recent studies have revealed that target eDNA concentrations are often higher in sediments (Turner et al. 2015; Kusanke et al. 2020; Nevers et al. 2020). Moreover, sediment sampling significantly enhances the probability of detecting benthic-dwelling diadromous fish, which predominantly inhabit the bottom of aquatic environments, such as the European eel (Burgoa Cardás et al. 2020c), the European flounder, or sea lamprey larvae which have extended development phases burrowed in sediments (Baltazar-Soares et al. 2022a).

However, it is worth noting that eDNA has been observed to persist for longer durations in sediments compared to DNA suspended or dissolved in water. This prolonged persistence is largely attributed to particle settling and/or the delayed degradation of DNA molecules adsorbed onto sediment (Turner et al. 2015). Consequently, while sediments offer extended detection windows, water is often considered a more accurate and reliable source of eDNA (Turner et al. 2015) advocating for its preference over sediment collection in eDNA analyses. To improve the likelihood of detecting benthic species, one strategy is to sample water close to the bottom. Notably, studies have reported higher detection rates for benthic species like the European eel in water samples collected from near-bottom locations (Burgoa Cardás et al. 2020c).

#### eDNA capture

In water-based eDNA studies, capturing eDNA prior to DNA extraction and assay execution is essential. While the optimal capture method may vary depending on environmental conditions and inhibitor concentrations (Kusanke et al. 2020; Troth et al. 2020), filtration has emerged as the primary choice for detecting diadromous fish (Fig. 5), especially recommended for rare species. However, when filtration is employed for eDNA capture, factors such as filter material and pore size can significantly influence recovered eDNA concentrations and subsequent target detection rates (Eichmiller et al. 2016; Hinlo et al. 2017; Troth et al. 2020). A closer examination of the employed volumes, filter materials, pore sizes, and filter preservation methods reveals a spectrum of options for diadromous fish species detection. However, certain procedures have emerged as predominantly used, including the use of 1L as the most adopted volume, filters composed of cellulose, with a pore size of 0.45 µm, and preservation at −20°C, before DNA extraction (Fig. 5).

**Fig. 5.**
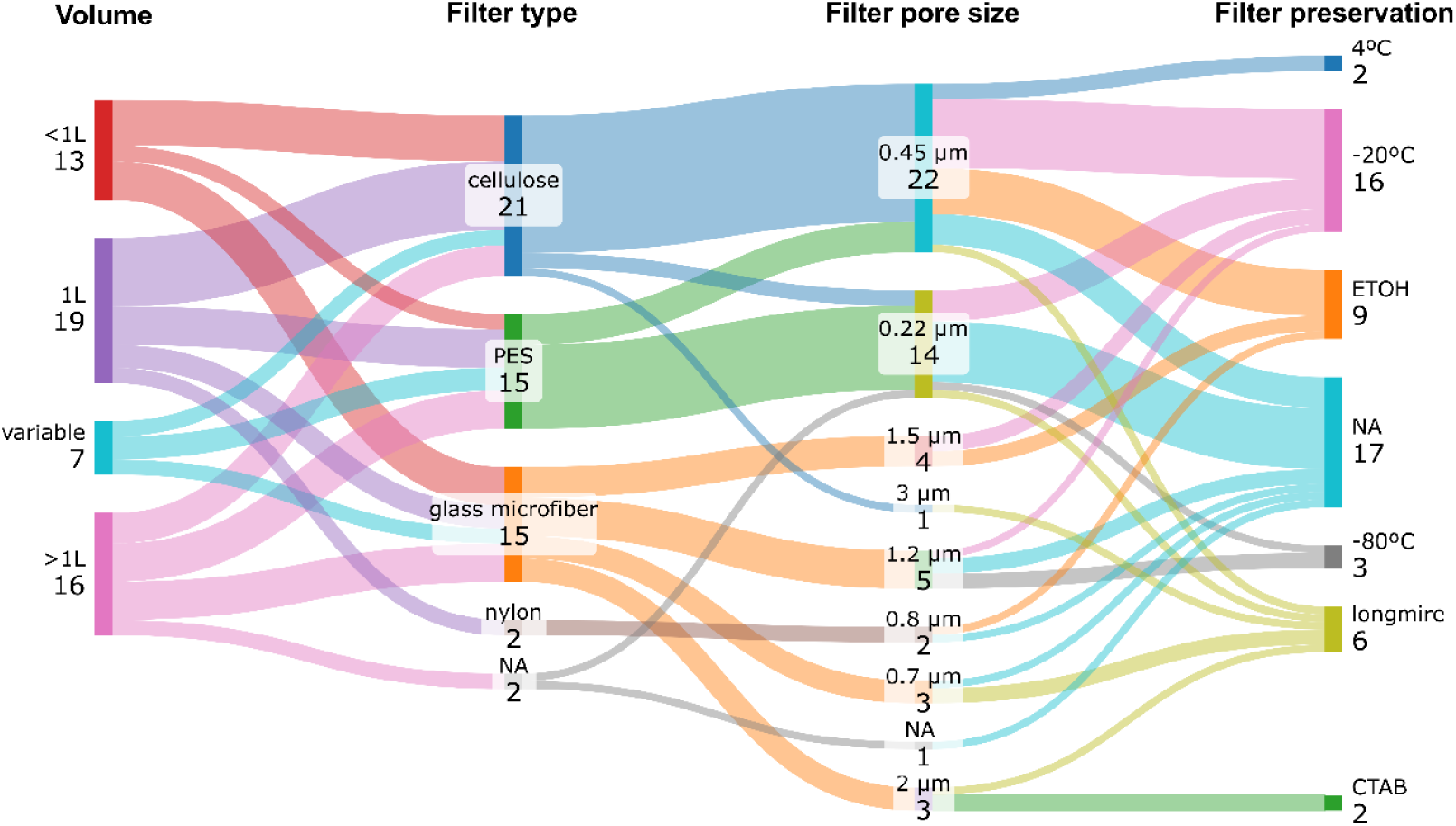
Sankey diagram presenting a synthesis of the predominant water volumes, types of filter material, pore sizes, and filter preservation methods commonly used in eDNA-based studies aimed at detecting diadromous fish species, with reported occurrence in Portugal.

While only a few studies have directly assessed the impact of these factors on eDNA-based detection, a comparison of cellulose nitrate (CN), mixed cellulose ester (MCE), polyethersulfone (PES), polycarbonate track-etched (PCTE), and glass fiber (GF) filters (all with similar pore sizes of 0.8 µm) revealed varied results. Although no concrete comparison has been found among our 49 analysed publications, a study by Hinlo and co-authors (2017) (Hinlo et al. 2017) found that CN filters generally yielded high eDNA and exhibited efficient detection of the oriental weather loach (*Misgurnus anguillicaudatus* (Cantor, 1842)). But the effectiveness of CN filters also appeared to depend on the specific combinations of DNA storage/preservation and DNA extraction methods employed.

It has been suggested that the varying binding affinities of DNA to different filter materials may be attributed to inherent filter properties. Generally, filters that retain particles both at the surface and within the filter’s matrices (such as GF, CN, and MCE filters) tend to yield greater DNA quantities compared to filters where particles are trapped only at the surface (i.e., retaining all particles larger than the pore size on the filter’s surface, such as PES and PCTE filters) (Eichmiller et al. 2016; Hinlo et al. 2017). Indeed, filtration of water samples through 1.2 μm GF and 0.45 μm MCE filters was more efficient in DNA recovery compared to using 0.22 μm enclosed PES filters, likely due to differences in DNA retention (Capo et al. 2019a).

Studies have demonstrated a negative correlation between captured eDNA and pore size, indicating that smaller pore sizes (ranging from 0.2 to 0.6 μm) offer greater sensitivity to changes in biomass, making them preferable for eDNA quantification (Minamoto et al. 2016; Kamoroff and Goldberg 2018). However, it is worth noting that most macro-organismal eDNA particles measure between 1 and 10 μm (Turner et al. 2014). Conversely, increasing the pore size can be advantageous when filtering larger water volumes or samples with high levels of suspended material, leading to higher eDNA yields (Eichmiller et al. 2016; Capo et al. 2019a).

The preservation method for filters has been shown to influence eDNA yields, especially when combined with specific DNA extraction methods (Hinlo et al. 2017). Hinlo and colleagues (Hinlo et al. 2017) found no significant difference in eDNA recovery between filters preserved in 95% ethanol or frozen at −20°C when using the Qiagen DNeasy kit for extraction. However, the PowerWater kit recovered significantly less eDNA from filters stored in ethanol, possibly due to compatibility issues. Storing filters in ethanol can be highly advantageous in field settings where access to ice is limited or electricity is unavailable, preventing immediate freezing of samples. This preservation method allows for transportation and storage at ambient temperatures (Minamoto et al. 2016; Hinlo et al. 2017) and has indeed emerged as the second most adopted preservation method for filters in studies employing eDNA for diadromous fish detection (Fig. 5).

#### eDNA extraction

The adoption of commercial kits, notably the DNeasy Blood & Tissue kit, has been favoured over non-commercial methods such as phenol–chloroform–isoamyl alcohol (PCI) or cetyltrimethylammonium bromide (CTAB)-based methods (Fig. 4). While comprehensive comparisons of DNA extraction protocols are lacking for diadromous fish species detection, several studies targeting other species have evaluated specific methods, particularly commercial kits, regarding their efficacy in extracting environmental DNA (eDNA) and on eDNA based detection (Piaggio et al. 2014; Eichmiller et al. 2016; Minamoto et al. 2016; Hinlo et al. 2017; Geerts et al. 2018; Lin et al. 2019; Kusanke et al. 2020; Steinmetz et al. 2021). For instance, Eichmiller and co-authors (2016) (Eichmiller et al. 2016) observed a trade-off between common carp eDNA yield and inhibitor removal, with kits featuring inhibitor removal steps yielding lower DNA quantities, but free from inhibitors (e.g., MoBio PowerSoil DNA Isolation Kit) compared to kits lacking these steps (e.g., MP Biomedicals FastDNA SPIN Kit and the Qiagen DNeasy Blood & Tissue Kit). Conversely, the combination of the Qiagen DNeasy Blood & Tissue kit with filtration using CN filters preserved in ethanol or stored at −20°C demonstrated superior performance in terms of cost and DNA recovery efficiency from the Oriental weather loach, when compared to other combinations (e.g., MoBio PowerWater either with ethanol or −20°C preserved filters) (Hinlo et al. 2017). The variances in cell lysis mechanisms -biochemical (e.g., DNeasy Blood & Tissue kit) *versus* mechanical (e.g., PowerWater) -likely account for the differing efficiencies of kits in releasing nucleic acids within the filter’s matrix (Hinlo et al. 2017).

In-house developed DNA extraction methods, such as phenol–chloroform–isoamyl alcohol (PCI) and cetyltrimethylammonium bromide (CTAB)-based methods, have been shown to be highly efficient and potentially more cost-effective than commercial kits (Geerts et al. 2018; Lin et al. 2019; Kusanke et al. 2020), but adopted less for eDNA-based detection of diadromous fish species (Fig. 4). For instance, the CTAB method exhibited significantly higher extraction efficiency compared to the PCI method or the Qiagen DNeasy Blood & Tissue kit, for eDNA-based active surveillance of the American bullfrog (Lin et al. 2019). Furthermore, an adapted PCI protocol demonstrated efficiency comparable to a commercial kit (MasterPure extraction kit) in crayfish DNA extraction from both tissue and filters (Geerts et al. 2018). However, it is worth noting that while PCI and CTAB methods are cost-effective, they require careful preparation and handling of toxic chemicals, and their protocols may demand more time compared to those utilizing commercial kits (Lin et al. 2019).

The selection of eDNA extraction methods is undoubtedly contingent upon research objectives and the chosen environments (e.g., freshwater *versus* marine; lentic *versus* lotic) (King et al. 2022). Additionally, factors such as personal preference, ease of use, and resource availability play significant roles in method selection.

#### Detection platform

Environmental DNA-based detection of diadromous fish species has been mainly achieved through 3 methodologies: quantitative real-time PCR (qPCR) (Gustavson et al. 2015; Carim et al. 2016; Jensen et al. 2018a; Atkinson et al. 2018a), digital droplet PCR (ddPCR) (Capo et al. 2019b, 2020; Thomson-Laing et al. 2023) and DNA metabarcoding (Harper et al. 2020; Macher et al. 2021; Barco et al. 2022; Halvorsen et al. 2023; Schallenberg et al. 2023) (Fig. 3b, Fig. 4, Supplementary Material 1, Table S2). The detection method adopted will greatly depend on the objective of the study. For instance, for more general biodiversity assessments, DNA metabarcoding would be advantageous. DNA metabarcoding employs broad-range primers designed to amplify the target gene across multiple species within a taxonomic group, such as metazoans or more specific groups such as fish, and use high-throughput sequencing (HTS) to sequence the resulting amplified DNA mixture (Taberlet et al. 2018). Bioinformatic tools are then used to process sequencing reads and to match them to genetic reference libraries. DNA metabarcoding, is, thus, often considered as the best qualitative tool suited for presence and absence of biodiversity surveys. Contrarily, qPCR, ddPCR and other single-species techniques apply assays using specific primers for targeting single or a very few species at a specific location and time, increasing the likelihood of detection. Targeted detection is particularly advantageous for “finding the needle in the haystack” and when the target is of high risk if it goes undetected (Morisette et al. 2020). In addition, qPCR and other single-species techniques can provide reliable measures of eDNA concentration (Schloesser 2018; Atkinson et al. 2018a). While the influence of the methodology on the detection of diadromous fish species has been relatively understudied, findings suggest that single species detection assays (i.e. ddPCR) exhibit higher sensitivity in detecting brown trout compared to DNA metabarcoding (Thomson-Laing et al. 2023). However, it is noteworthy that metabarcoding allows for the detection of additional fish species (Thomson-Laing et al. 2023). An illustrative example lies in the monitoring of non-native fish species using environmental DNA (eDNA). Combining both active and passive surveillance methodologies with eDNA samples may present significant advantages, especially in the context of non-native species management (Simmons et al. 2016).

Additionally, alternative methods for detecting diadromous fish species have emerged, including CRISPR-Cas technology (Williams et al. 2019, 2021, 2023), high-resolution melt curve (HRM) analysis (Minett et al. 2021b), and nested PCR-restriction fragment length polymorphism (PCR-RFLP) (Clusa et al. 2017; Fernandez et al. 2018a). Advantages of these methods over qPCR or ddPCR may include high precision and reduced risk of off-target effects with CRISPR-Cas, rapid detection capabilities, and the elimination of the need for additional probes or labelled primers with HRM or PCR-RFLP, which significantly reduces associated costs.

### Emerging applications of eDNA in monitoring and conservation of diadromous fish species in Portugal

While most of the studies analysed here focus primarily on the development and validation of eDNA-based assays for detecting diadromous fish species, a few have employed eDNA for assessing diadromous fish distribution (primarily relying on metabarcoding, accounting for approximately 26% of the studies, as shown in Supplementary Material 1, Table S2) (Fernandez et al. 2018b; Griffiths et al. 2020; Consuegra et al. 2021b; Barco et al. 2022). Although this highlights a gap in the literature - out of the 49 studies reviewed, only a small percentage focused on understanding the distribution of diadromous fish species, while none have, for example, addressed diadromous fish species diets - practical applications have emerged in recent years and could be implemented in the future in Portugal:

#### Track changes in diadromous fish species populations and identify critical spawning and nursery areas

Rapid eDNA analysis from many aquatic ecosystems may help focus traditional assessment efforts, thereby improving species detection efficiency over time and providing data on population trends, distribution, and spawning success. It can also help to identify previously unknown habitats that are important for the species. For instance, the presence of eel has been effectively detected through an eDNA-based qPCR assay in the Adriatic Sea and in the complex Dinaric karst freshwater ecosystem (Vucić et al. 2023b), and this information can be used to identify important conservation areas. The use of eDNA metabarcoding to assess fish species diversity in marine protected areas of the North Sea allowed documentation of three additional species to those found in the same year with trawls, including the critically endangered European eel (Barco et al. 2022). eDNA-based qPCR specific assays for shads (Antognazza et al. 2019, 2021) and for sea lamprey (Bracken et al. 2019) have been used to observe the timing and spatial range of spawning migrations in rivers from the UK and, thus, to identify critical nursery areas that need protection or restoration. In future studies, eDNA could be combined with other techniques to enhance the understanding of migration patterns and the identification of reproduction or maternity areas. For instance, acoustic sensors can be employed to track the movement of individuals or groups along migration routes, while eDNA analysis can be used to confirm the presence of specific species in key areas. This integrated approach could provide a more comprehensive picture of habitat use and connectivity (see below), enabling the identification of critical breeding or nursery zones. Combining methodologies like telemetry, acoustic tracking, and eDNA (Harris et al. 2022; Dunn et al. 2023) could also help address challenges such as species-specific detection, seasonal variations, and the influence of environmental factors on migration and reproduction.

#### Evaluate the impact of barriers and monitor the success of habitat restoration projects

eDNA monitoring can be also valuable in assessing the impacts of migration barriers like dams on anadromous fish species movement and spawning (Halvorsen et al. 2020; Consuegra et al. 2021a). An eDNA-based qPCR specific assay for detecting European eel through eDNA in rivers from Norway was used to evaluate how hydroelectric power stations affect their migration patterns (Halvorsen et al. 2020). The size of barriers was also evaluated on species such as Atlantic salmon, brown trout, shads, and lamprey in River Allier (France), using eDNA metabarcoding (Consuegra et al. 2021a). This information can inform decisions on barrier removal or the installation of fish passage solutions. eDNA metabarcoding proved to be highly sensitive to monitor the effects of barrier removal in fish community composition, including diadromous fish species such as the European eel, Atlantic salmon, and brown trout in the UK rivers (Muha et al. 2021). This information was critical for assessing the effectiveness of conservation measures, by providing evidence of species recolonization and use of restored areas. Once again, eDNA can be a complement to telemetry, acoustic or visual techniques to track the movement of individuals, while also confirming the presence of specific species in key areas.

#### Trophic interactions and predator-prey dynamics

In addition to species detection and habitat monitoring, eDNA analysis has also proven valuable in identifying predators of diadromous fish species. By analyzing the stomach contents of predators, eDNA can provide precise identification of prey species, including diadromous fish, which are often difficult to recognize through traditional visual methods. This offers new insights into predator-prey relationships and helps assess the ecological pressures faced by these species, aiding in the development of more effective conservation strategies. For example, an eDNA-qPCR specific assay was used to detect European eel in potential predators’ stomach contents in the Sargasso Sea (Jensen et al. 2018a). The detection of species in almost 10% of all fish stomachs investigated, representing 6 species, gave important information on how mesopelagic fishes in the Sargasso Sea can predate to some extent eel larvae. For instance, eDNA metabarcoding has been used to investigate the diet and niche partitioning between native European otters and invasive American mink, indicating that diadromous fish species such as European flounder, European eel and Brown lamprey make up 80% of otters diet, while minks’ diet was dominated by terrestrial birds and mammals with a minimum overlap between the two species (Harper et al. 2020). Although not extensively explored in the reviewed literature, eDNA-based tools hold significant potential for advancing our understanding of species’ diets across various life stages, from juveniles to adults. For example, DNA barcoding has previously demonstrated its effectiveness in providing a qualitative assessment of the diet of European eel larvae in the Sargasso Sea, even identifying highly unrecognizable prey items (Riemann et al. 2010). Building on this, DNA metabarcoding and eDNA offer even greater potential to identify a broader range of species present in the diet, delivering deeper insights into trophic interactions and ecological dynamics. While some studies in Portugal have employed stable isotopes (França et al. 2011; Dias et al. 2019), eDNA presents distinct advantages by yielding more detailed and comprehensive data. This approach can address existing uncertainties, particularly in situations where traditional methods, such as stomach content analysis, face limitations. These include challenges posed by the small size of prey, incomplete or difficult-to-identify organisms, or the elusive behaviour of juvenile individuals. In fact, for certain life stages, such as juveniles (Dias et al. 2019) and even adults, dietary information is often sparse or entirely absent. eDNA, therefore, holds great promise for filling these knowledge gaps and significantly expanding research opportunities in this field.

#### Biomass estimation

Although the relationship between eDNA concentration and biomass is not always straightforward (Knudsen et al. 2019b) and can vary depending on several factors (e.g. water flow, temperature, DNA degradation rates, and the species’ behaviour), eDNA can indeed be used as an indirect measure of species biomass, if specific assays are developed for a particular species. For instance, by applying correction for altitude and a linear model, Fernandez and co-authors (2023) (Fernandez et al. 2023) were able to predict eel biomass by using an eel-specific qPCR marker in different rivers of northern Spain. The method was validated by electrofishing surveys. In addition, a strong positive correlation between adult sea lamprey densities of 2, 20, and 200 individuals/2000L and eDNA concentrations was found in laboratory tanks using eDNA-based specific assays (Schloesser 2018). The same can be applied to breeding sites to explore whether locations with a higher aggregation of adults also have more eggs and more DNA. This could be particularly interesting to investigate in both water and sediment, as the differences in species ecologies may influence DNA distribution, persistence, and detectability across these distinct matrices. Other factors should be also taken into account such as the proximity of the sampled patch to the presence of individuals (Baltazar-Soares et al. 2022b).

### Advances and challenges of eDNA-based monitoring for diadromous fish conservation in Portugal

There are several advantages to using eDNA-based monitoring for detecting diadromous fish, underscoring its potential to encourage the adoption of these tools to enhance knowledge and conservation of these species in Portugal: 1) eDNA involves collecting water (Gustavson et al. 2015; Antognazza et al. 2019; Capo et al. 2019b) or sediment samples (Baltazar-Soares et al. 2022a) to detect the presence of species without needing to capture or disturb the organisms. This is particularly beneficial for several diadromous fish species (e.g., sea lamprey, European eel, European flounder), which may be elusive or inhabit difficult-to-sample sites; 2) eDNA can detect low-abundance species or early life stages that might be missed by traditional survey methods.

This sensitivity is crucial for monitoring fish populations, especially in the early stages of their spawning migration (Baltazar-Soares et al. 2022a) or species that are becoming rarer (e.g., sea lamprey); 3) sampling can be more cost-effective and less labour-intensive than traditional methods like electrofishing or netting. It reduces the need for extensive fieldwork and specialized equipment (Rusch et al. 2018; Fernández et al. 2019b; Burgoa Cardás et al. 2020c; Jacobsen et al. 2023); 4) eDNA analysis can provide quicker results compared to some traditional monitoring techniques. This allows for more timely decision-making and adaptive management strategies (Burgoa Cardás et al. 2020c; Vucić et al. 2023b), and, finally, 5) eDNA can be used to monitor multiple locations simultaneously, providing a comprehensive view of diadromous fish distribution across large areas (Knudsen et al. 2019a; Barco et al. 2022; Vucić et al. 2023b) and different river systems (Fernandez et al. 2018b; Hallam et al. 2021).

There is no doubt that eDNA-based monitoring can greatly improve the conservation, management, and restoration of diadromous fish species populations across various Portuguese aquatic ecosystems. However, there are still several challenges and considerations on the use of eDNA-based detection for monitoring diadromous fish species, such as the dispersion and dilution of eDNA in large lotic systems (Laporte et al. 2020b, c; Wood et al. 2020) or degradation over time, potentially complicating the interpretation of results regarding the exact location of the species. Various environmental factors, such as river flow (Malekian et al. 2018), temperature (Robson et al. 2016; Jo et al. 2020; Caza-Allard et al. 2022), pH (Tsuji et al. 2017), among others can also affect eDNA detection rates (Duarte et al. 2023) and that need to be accounted in study designs.

In addition, the use of species-specific eDNA assays that can reliably distinguish diadromous fish species from other similar species is crucial for accurate monitoring. For that, DNA sequences from both target, closely related and co-occurring species is crucial to test the specificity of the assays. Lack of sufficient specificity can lead to both false positives and false negatives, especially when abundant, closely related species are present (Wilcox et al. 2014). Moreover, closely related species may hybridize (Antognazza et al. 2019, 2021), further complicating the design of specific assays capable of distinguishing hybrids from non-hybrids due to the maternal inheritance of mitochondrial DNA. For example, the assay developed for shads (Antognazza et al. 2019, 2021) cannot differentiate between allis shad and twaite shad using the COI region, highlighting the need to explore other genetic markers that can reliably distinguish between the two species. Additionally, distinguishing between different populations or ecotypes of the same species that co-exist in the same habitat, such as *Salmo trutta*, can be challenging. Resident populations (brown trout) are often difficult to differentiate genetically from migratory sea trout populations (for example no differences have been identified in COI sequences – Supplementary Material 1, Table S3). However, exploring other genetic markers or employing tools that can assess differences at the population level (e.g., microsatellites) (Aurelle and Berrebi 1998) may help address this issue.

Assays previously developed for species detection at a specific geographic location may be ineffective in other regions due to sequence matches with co-occurring species (Ogata et al. 2022) or intraspecific variation (Oficialdegui et al. 2019). These case studies highlight the need to refine eDNA-based assays for target species by considering site-specific genotypes. A quick look into the Barcode of life Data System database (BOLD) shows that all diadromous fish species found in Portugal are represented by COI sequences, but not all species are represented by sequences derived from Portuguese specimens. For example, sequences for *Lampetra fluviatilis*, *Salmo salar*, and *Salmo trutta* in the database were generated for specimens collected in other countries. In addition, Portuguese specimens are represented by a very low number of sequences (≤5 sequences: *Alosa alosa,* 1 sequence; *A. fallax*, 1 sequence; *Anguilla anguilla*, 1 sequence; *Chelon ramada*, 5 sequences; *Petromyzon marinus*, 2 sequences and *Platichthys flesus*, 4 sequences) (Supplementary Material 1, Table S4). This lack of region-specific reference sequences can lead to decreased accuracy in species identification and monitoring efforts, particularly when local genetic variants differ from those in other geographic regions, which can be particularly relevant for species with homing behavior, such as salmonids and clupeids (Ogata et al. 2022). This creates strong genetic structuring within species because populations become adapted to their local environments, which is why region-specific reference sequences are so important when using eDNA for species identification. Such gaps in the reference libraries may hinder the effectiveness of eDNA assays, especially in areas where species exhibit cryptic diversity or where population structure is significant (Oficialdegui et al. 2019). Expanding these reference datasets will facilitate more accurate species detection and allow for better-informed conservation and management strategies. To enhance the reliability of eDNA-based monitoring for diadromous fish species in Portugal, it is crucial to expand the representation of local population sequences in reference databases like BOLD. In addition, existing assays should be rigorously tested through a combination of *in silico*, *in vitro*, and *in vivo* experiments to evaluate their performance. Where necessary, new assays may need to be developed to ensure accurate species detection, reflecting the unique genetic diversity of local populations.

End-user confidence in adopting eDNA-based monitoring for diadromous fish species in Portugal will depend on several factors, including the validation and standardization of methods to ensure consistent, accurate results across regions and labs, which builds trust in the technology (Darling 2019; Mosher et al. 2020). Establishing standardized protocols for eDNA sampling, analysis, and data interpretation is also essential to ensure consistency and reliability of results, and based in this review analysis, at least in what respects the experimental workflow these have been widely variable, which also difficult comparisons among studies. Simplified workflows, user-friendly tools, and clear result interpretation can make eDNA monitoring accessible, while case studies in other regions that showed tangible conservation outcomes, such as improved habitat protection or population insights, reinforce its value (Fernandez et al. 2018b; Consuegra et al. 2021b; Vucić et al. 2023b). Educational and collaborative initiatives further increase familiarity and trust, allowing end-users to engage directly with the methodology. Here, collaborations with fishers could even be established to collect samples or to provide information about migration routes, spawning areas, among other relevant information. Finally, regulatory support from national or regional bodies would solidify eDNA’s legitimacy, encouraging wider adoption in Portugal’s conservation and fishery monitoring practices.

**Fig. 6.**
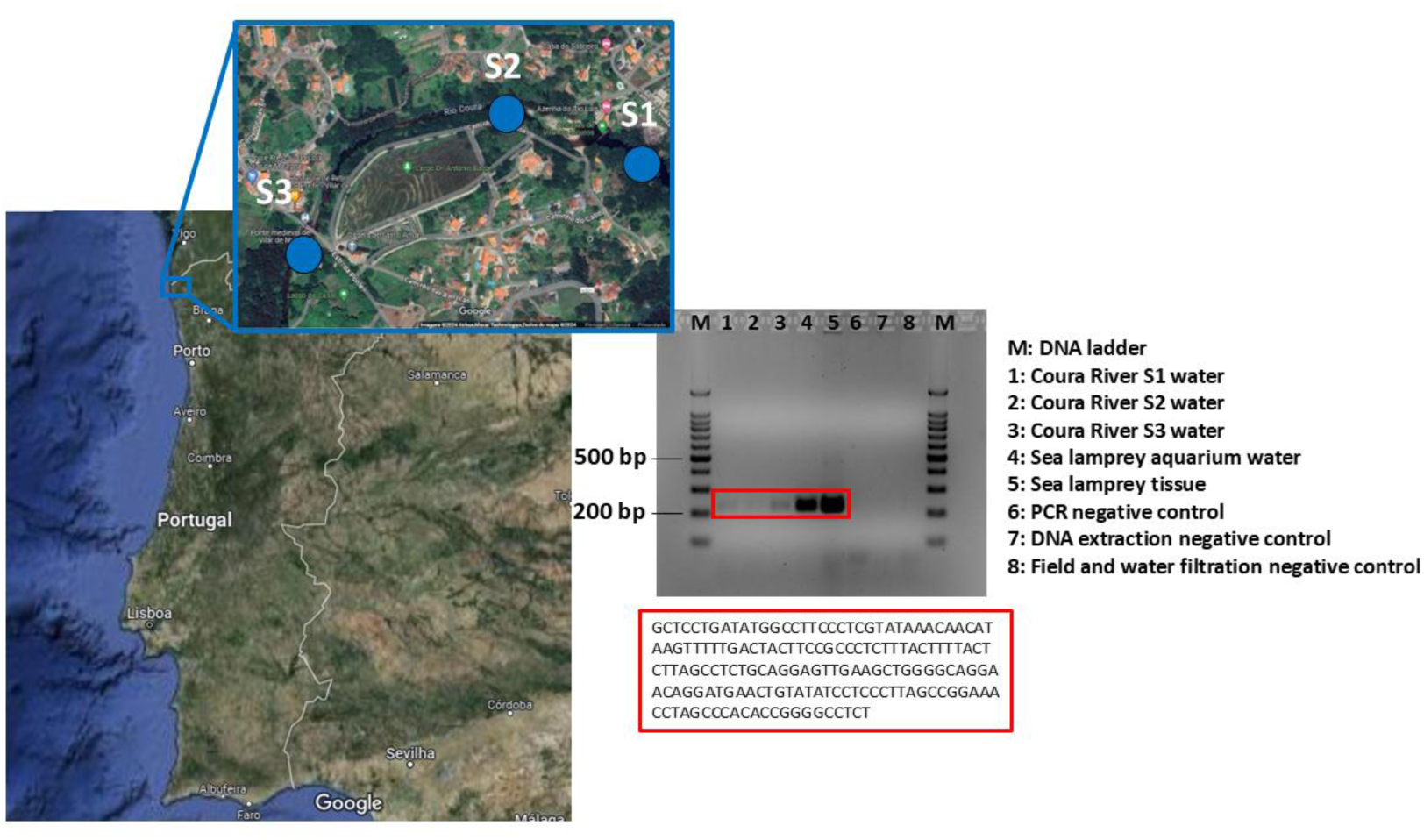
Agarose gel electrophoresis of PCR products using the species-specific primers for the detection of sea lamprey at three sites of the Coura River, in water samples collected in the Aquamuseum aquariums with sea lamprey and in sea lamprey tissue. Source of the map: Google maps.

Additionally, high-throughput Illumina MiSeq sequencing with general COI primers (Lobo et al. 2013) was used to evaluate the broader detection potential. Although more targeted markers like 12S or 16S have been more adopted for the detection of fish species through eDNA metabarcoding (Cawthorn et al. 2012; Laporte et al. 2020b; Barco et al. 2022), since it could reduce non-target amplification, they often have less comprehensive databases and lower species-level resolution (Cawthorn et al. 2012). In addition, the choice of COI allowed direct comparison with the species-specific assay employed for this marker, providing a valuable opportunity to evaluate the performance of both approaches. Illumina sequencing generated 253,438 raw reads, which were filtered to 167,561 sequences (after removal of short length reads, de-multiplexing, primer removal, de-replication, and chimera removal) (Supplementary Material 1, Table S6). Taxonomic matches exceeding 97% were highest in Aquamuseum samples (13%), where 63% were attributed to *P. marinus*. The highest species richness was observed in Coura River sample S1 (13 taxa), while Aquamuseum samples contained fewer taxa, reflecting the controlled aquarium environment (Supplementary Material 1, Table S7). These findings underscore the advantages of DNA metabarcoding over targeted detection, as it enables simultaneous detection of the target species (*P. marinus*) and other taxa, including the common non-native *Gobio lozanoi*, the rare non-native *Oncorhynchus mykiss*, and the elusive native amphibian species *Salamandra salamandra* and *Triturus marmoratus*. Although this preliminary test demonstrates the high potential of eDNA for detecting sea lamprey, further validation is essential, including tests at sites where sea lamprey is not expected to occur.

## Author contributions

F.O.C. and S.D. conceptualized the study. S.D. made the literature review, analyzed the data, prepared the figures and wrote the main manuscript. A.M.S., C.A., R.S., and S.D. carried out the experimental work for the pilot study. F.O.C. and S.D. secured the funding. All authors reviewed and approved the final manuscript.

## Acknowledgments

This work was funded by the project “River2Ocean – Socio-ecological and biotechnological solutions for the conservation and valorization of aquatic biodiversity in the Minho Region” (NORTE-01-0145-FEDER-000068), co-financed by the European Regional Development Fund (ERDF), through Programa Operacional Regional do Norte (NORTE 2020) and by the “Contrato-Programa” UIDB/04050/2020, funded by national funds through the Foundation for Science and Technology (FCT I.P). Financial support granted by the FCT to SD (https://doi.org/10.54499/CEECIND/00667/2017/CP1458/CT0001), is also acknowledged.

