## Supplementary material 2 for "Detection of diadromous fish using environmental DNA: prospects for its use in conservation of endangered species occurring in Portugal"

### Materials and Methods

*Detection of sea lamprey through eDNA in the Coura River (North of Portugal)*

To validate and assess the efficacy of previously developed primers for sea lamprey detection (Gingera et al. 2016), three distinct assays were conducted: 1) *in vitro* testing, using DNA extracted from sea lamprey tissue (Peixaria S. João, Esposende, Portugal) and 2) *in situ* testing (aquarium testing) where the primers were tested against eDNA extracted from water samples collected from aquaria housing sea lamprey specimens. These aquaria were located at the Minho River Aquamuseu (https://aquamuseu.cm-vncerveira.pt/) and 3) *in situ* testing (natural habitat) where the primers were further evaluated against eDNA extracted from water samples collected from sites along the Coura River (41.889369, -8.786620), where the presence of sea lamprey has been documented (Sousa et al. 2012). For each water-based assay, 3 replicates of 1 L each were used, with samples filtered through 0.45 µm pore size nitrocellulose filters (Millipore).

DNA extraction from both sea lamprey tissue and filters was conducted using the DNeasy Blood & Tissue kit (Qiagen) following the manufacturer's instructions. Each PCR reaction had a total volume of 25 µL, comprising 1× Speedy Supreme NZYTaq 2x colourless master mix (NZytech), 0.2 µM of each primer (Gingera et al. 2016), and 8 µL of DNA extract.

PCR program consisted of an initial 5-minute denaturation step at 95 ºC; followed by 35 cycles of denaturation at 95 ºC for 30 seconds, annealing at 55 ºC for 30 seconds, and elongation at 72 ºC for 30 seconds; with a final elongation step at 72 ºC for 5 minutes, according to Gingera and co-authors (Gingera et al., 2016).

The amplified products were visualized through electrophoresis on a 2% agarose gel stained with GreenSafe (NZytech). Positive amplifications were subsequently confirmed by Sanger sequencing of the amplicons at Macrogen Europe. Additionally, DNA extracts underwent high-throughput sequencing on an Illumina Miseq platform (Genoinseq, Cantanhede, Portugal).

Sample preparation for Illumina Sequencing involved COI gene amplification. The DNA underwent initial amplification of the region using specific primers. Subsequently, a limited-cycle PCR reaction was conducted to append sequencing adapters and dual indexes.

The first PCR reactions were carried out for each sample utilizing the KAPA HiFi HotStart PCR Kit (Kapabiosystems, Cape Town, South Africa) as per the manufacturer's recommendations. The reaction mixture contained 0.3 µM of each PCR primer (forward primer mICOIintF 5'-GGWACWGGWTGAACWGTWTAYCCYCC-3' and reverse primer LoboR1 5'–TAAACYTCWGGRTGWCCRAARAAYCA -3') (Leray et al. 2013; Lobo et al. 2013) and 5 µL of template DNA in a total volume of 25 µL.

PCR conditions comprised a 3-minute denaturation at 95 ºC, followed by 35 cycles of 98 ºC for 20 s, 60 ºC for 30 s, and 72 ºC for 30 s, with a final extension at 72 ºC for 5 min. Negative PCR controls were included for all amplification steps.

In the second PCR reactions, indexes and sequencing adapters were added to both ends of the amplified target region according to the manufacturer's recommendations (Illumina 2013). PCR products were then purified and normalized using the SequalPrep 96-well plate kit (ThermoFisher Scientific, Waltham, USA), pooled, and pair-end sequenced on the Illumina MiSeq® sequencer with the MiSeq reagent Kit v3 (600 cycles), following the manufacturer's instructions (Illumina, San Diego, CA, USA) at Genoinseq (Cantanhede, Portugal). Sequence data processing was conducted at Genoinseq (Cantanhede, Portugal).

Raw reads were extracted from the Illumina MiSeq® System in fastq format and quality-filtered using PRINSEQ version 0.20.4 (Schmieder and Edwards 2011) to eliminate sequencing adapters, trim bases with an average quality lower than Q25 in a 5-base window, and remove reads shorter than 150 bases. Additionally, bases with an average quality lower than Q25 in a 5-base window were trimmed. The filtered forward and reverse reads provided by the sequencing facility were merged by overlapping paired-end reads in mothur (make.contigs function, default alignment) (version 1.39.5), with primer sequences also removed (trim.seqs function, default) (Schloss et al. 2009; Kozich et al. 2013). The processed reads were then analyzed in mBrave – the Multiplex Barcode Research and Visualization Environment (www.mbrave.net) (Ratnasingham 2019), which is integrated with BOLD (Ratnasingham and Hebert 2007). Raw sequencing data of each dataset was deposited in the Sequence Read Archive (SRA) of NCBI in the BioProject ID PRJNA1232697 (https://www.ncbi.nlm.nih.gov/bioproject/1232697).
